# qEEG Analysis in the Diagnosis of Alzheimer’s Disease; a Comparison of Functional Connectivity and Spectral Analysis

**DOI:** 10.1101/2022.01.10.475756

**Authors:** Maria Semeli Frangopoulou, Maryam Alimardani

## Abstract

Alzheimer’s disease (AD) is a brain disorder that is mainly characterized by a progressive degeneration of neurons in the brain, causing a decline in cognitive abilities and difficulties in engaging in day-to-day activities. This study compares an FFT-based spectral analysis against a functional connectivity analysis based on phase synchronization, for finding known differences between AD patients and Healthy Control (HC) subjects. Both of these quantitative analysis methods were applied on a dataset comprising bipolar EEG montages’ values from 20 diagnosed AD patients and 20 age-matched HC subjects. Additionally, an attempt was made to localize the identified AD-induced brain activity effects in AD patients. The obtained results showed the advantage of the functional connectivity analysis method compared to a simple spectral analysis. Specifically, while spectral analysis could not find any significant differences between the AD and HC groups, the functional connectivity analysis showed statistically higher synchronization levels in the AD group in the lower frequency bands (delta and theta), suggesting that the AD patients’ brains are in a ‘phase-locked’ state. Further comparison of functional connectivity between the homotopic regions confirmed that the traits of AD were localized in the centro-parietal and centro-temporal areas in the theta frequency band (4-8 Hz). The contribution of this study is that it applies a neural metric for Alzheimer’s detection from a data science perspective rather than from a neuroscience one. The study shows that the combination of bipolar derivations with phase synchronization yields similar results to comparable studies employing alternative analysis methods.

## Introduction

Alzheimer’s disease (AD) is a brain disorder that is mainly characterized by a progressive degeneration of neurons in the brain. As the disease progresses, a cortical disconnection occurs, causing a deficit in memory and a decline in other cognitive capabilities [1]. AD-related effects on the patient’s brain can be identified with various tools, one option being the electroencephalogram (EEG), which measures the electrical activity of the brain. EEG is a fast and non-invasive method that provides a high temporal resolution. However, it lacks in spatial resolution, meaning that it is not the most precise method for the diagnosis of a brain disorder.

Quantitative EEG (qEEG) analysis takes EEG recordings, commonly interpreted by clinicians using visualization tools, one step further, giving the possibility of digitally processing and presenting the signal characteristics in spectral and spatial domains [2]. In a spectral analysis, a given signal is broken down and examined in the frequency domain. This type of analysis is useful when finding differences between patients diagnosed with a disorder and healthy individuals, by examining relevant frequency bands to identify a noticeable change in the activity within a particular frequency band [3]. A very common yet powerful tool used in spectral analysis is the Fast Fourier Transformation (FFT) [4]. This algorithm can be used to find band-specific differences by calculating the power of each band separately.

When conducting a spectral analysis, the power spectral density (PSD) is often used to determine differences in brain activity between frequency bands. Previous studies have shown that compared to healthy controls (HC), AD patients show an increase of PSD in the theta band and a decrease in the alpha band [3, 5, 6]. In AD diagnosis specifically, a spectral analysis can show discrepancies between AD and other types of dementia, such as vascular dementia (VaD) [1].

However, while these studies suggest that EEG spectral analysis may differentiate AD patients from HC [7], several other studies that examined the process of AD have concluded that this brain disorder is involved with changes in the distributed networks related to memory [8] and that the changes observed in the frequency bands may not sufficiently reflect this. Moreover, as mentioned above, patients suffering from AD experience a cortical disconnection. It is therefore important to examine various Regions of Interest (ROIs) that are affected by the disease. Hence, more reliable signal processing methods are required to capture the complexity of this disorder and investigate the processes that underlie the occurring symptoms [9, 10]. An alternative to a spectral analysis is the connectivity analysis; a method which allows to study the communications between different regions of the brain [10].

Functional connectivity analysis measures the degree of synchronization between two EEG signals; a higher connectivity indicates more effective communication between the examined brain regions [11]. There are several ways of conducting a functional connectivity analysis. For instance, **Coherence analysis** has been used exhaustively in detecting differences between AD patients and HC. Recent studies indicate a decrease in the coherence levels between ROIs for the AD [3, 12]. Although coherence has brought some novelty in studies involving AD patients, it is worth mentioning that it solely takes into account linear correlations, thus not considering nonlinear interactions.

Nonlinear correlations, on the other hand, can give crucial information in a functional connectivity analysis. A widely used method for this is the **phase synchronization (PS)** analysis. PS looks at the oscillatory activity in two brain regions in terms of their phases [13]. The oscillations are therefore said to be synchronized if their phases are similar. PS excels over coherence analysis in terms of being able to account for nonlinearity [14]. Moreover, a study has shown that differences have been found in terms of synchronization between within-band connections and between-band connections (e.g., within delta band; between delta and theta bands) [15]. This study in particular also discovered that AD patients showed much lower strength of synchronization for between-frequency band analysis when compared to HC.

PS has several indices of measurement, with the **phase-lag index (PLI)** and **phase-locking value (PLV)** being the most used measures [16]. The PLI gets a time-series of phase differences and computes the asymmetry corresponding to the distribution of these phase differences [17]. In a recent PS study using the PLI as the index of choice, results showed that in AD patients, the lower alpha band presented a decrease in functional connectivity situated in the posterior region [18]. On the other hand, PLV looks at the consistency in phase difference. The PLV value ranges from 0, indicating random phase differences, to 1 indicating a fixed phase difference [19]. For example, a study performing cross-frequency coupling (CFC) using PLV on AD patients reached the conclusion that, oscillations in the alpha band, and more specifically around the dominant peak, are phase-locked with the gamma band power [20]. Results were observable in the posterior region of the brain suggesting that AD elicits a region-specific change in functional connectivity.

In sum, the current state-of-the-art calls for a comparison between computational methods that are used for diagnosis of Alzheimer’s disease. So far, most studies have reported the outcomes of either a spectral analysis or a connectivity analysis [6, 21–23]. However, conducting a connectivity analysis and comparing it with a spectral analysis using the same dataset presents two advantages; 1) it shows which method can yield the most accurate and complete information in AD diagnosis [3, 24], and 2) it can identify the affected ROIs instead of solely looking at whether the patient suffers from AD. By finding potential ROIs, it is believed that this technique could help predicting AD in its early stages of development [10].

The proposed study serves as a comparison between the two methods, namely the spectral and connectivity analyses. The two types of analysis were conducted on a set of EEG recordings obtained from patients suffering from AD and from their respective healthy controls (HC), in an attempt to address the following research question:

*RQ1: How does a functional connectivity analysis perform against a spectral analysis in finding differences between patients diagnosed with Alzheimer’s disease (AD) and healthy controls (HC)?*

Moreover, this study attempted to answer a secondary research question:

*RQ2: Can a functional connectivity analysis localize the differences identified in the brain activity of AD subjects when compared to that of the HCs?*

To answer this question, a series of statistical tests were made using the results provided by the connectivity analysis.

## Methods and Materials

### Dataset and Preprocessing

The EEG dataset was provided by the University of Sheffield under a relevant NDA. All subjects were informed about the experiment and signed an informed consent form. The dataset consists of 12-seconds, eyes-open recordings of 20 AD-diagnosed patients and 20 age-matched HC, younger than 70 years of age (Table 1).

**Table 1:** General information of the AD and HC groups including sample size, age mean with standard deviation and gender ratio per group

|  | AD | HC |
| --- | --- | --- |
| Size | $N = 20$ | $N = 20$ |
| Age | 60 ($SD = 4.40$) | 61 ($SD = 6.67$) |
| Gender (F/M) | 8/12 | 12/8 |

The participants’ EEGs were recorded using the International 10-20 system [25]. To reduce volume conduction effects from a common reference [26], 23 bipolar derivations were used in this study. Figure 1 gives an overview of the electrodes and bipolar channels. More specifically, the following bipolar channels were used: F8-F4, F7-F3, F4-C4, F3-C3, F4-FZ, FZ-CZ, F3-FZ, T4-C4, T3-C3, C4-CZ, C3-CZ, CZ-PZ, C4-P4, C3-P3, T4-T6, T3-T5, P4-PZ, P3-PZ, T6-O2, T5-O1, P4-O2, P3-O1, O2-O1. These bipolar channels are the most commonly used in clinical practice [27]. During the recording, the participants were instructed to reduce their movements and not to think of anything in particular (i.e., resting state EEG).

**Figure 1:**
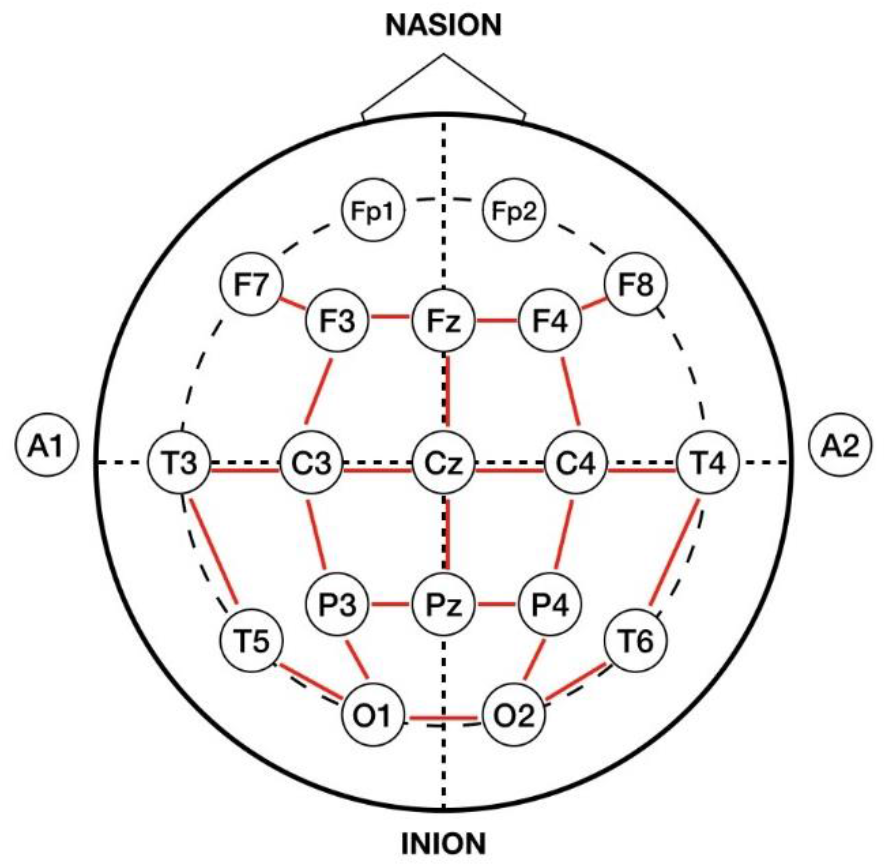
EEG signals were collected from 23 bipolar channels based on the 10-20 international system

The raw EEG signals were preprocessed in EEGLAB (v.2021.0), MATLAB. First signals were downsampled to 500Hz. Next, a band-pass filter was applied between 0.1 and 100 Hz using EEGLAB functions following the requirements used for the phase synchronization (see Section ‘Functional Connectivity Analysis’) to avoid phase distortion. Additionally, a notch filter was used to attenuate signals in 48-52 Hz.

### Spectral Analysis

The power spectral density (PSD) of the entire signal for each of the bipolar montages was calculated using EEGLAB’s spectopo() function. This function makes use of the FFT algorithm to extract and plot the PSD. The signal was subsequently divided into five frequency bands: delta (1-4 Hz), theta (4-8 Hz), alpha (8-13 Hz), beta (13-30 Hz), gamma (36-44 Hz) and the mean power in each band was computed. These ranges were selected according to [28] and were also used in the connectivity analysis. A Shapiro-Wilk test was applied to the data to check for normality and subsequently, a Mann-Whitney U-test was used to compare band power medians between the groups AD and HC.

### Functional Connectivity Analysis

Functional connectivity analysis was carried out using the PLV index [28]. First, a continuous wavelet transform was applied (i.e., the Complex Morlet wavelet), with this wavelet being used as a kernel to compute the PLV, which is defined by Equation 1:

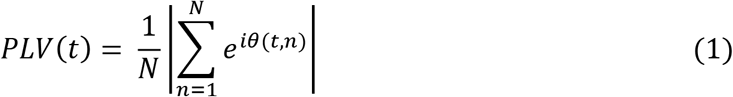

where *n* is an index for the trial number and *θ* indicates the phase difference. The phase-locking value yielded by PLV ranges from 0 to 1, with 1 indicating that two signals have an identical relative phase across N trials. Conversely, values that approach 0 indicate little to no phase synchrony between the signals. For every subject, the PLV was calculated for all possible 253 bipolar channel combinations in five frequency bands as defined above. Next, inspired by [29], ‘Global Connectivity’ and ‘Homotopic Pair Connectivity’ were computed using the extracted PLV values and were compared between the groups.

#### Global Connectivity

Global Connectivity was computed by averaging all 253 PLV values that were obtained per frequency band. This led to a total of five PLV_mean_ values per subject (i.e., one PLV_mean_ per frequency band). Following the Shapiro-Wilk test, a Mann-Whitney test was used to compare the mean PLVs between the AD and HC groups. The aim of this evaluation was to determine whether band-specific differences could be found in the global functional connectivity of the AD subjects against the HCs.

#### Homotopic Pair Connectivity

Homotopic Pair Connectivity was computed by focusing on certain pairs of bipolar derivations that were homotopic in the Left and Right brain hemispheres (mirror areas of the brain hemispheres). Based on previous classifications [30, 31], four pairs that were, in part, shown most affected by Alzheimer’s disease were selected. These pairs are demonstrated in Figure 2. Pair A consisted of the homotopic pair located in the centro-parietal region of the brain (C3-P3 & C4-P4). Pair B corresponded to the pair in the fronto-central area (F3-C3 & F4-C4), Pair C consisted of electrodes located in the parieto-occipital region (P3-O1 & P4-O2) and Pair D consisted of electrodes placed in the centro-temporal area (C3-T3 & C4-T4). For each pair, the PLV was computed in the five frequency bands and a Mann Whitney U-test was carried out to compare the band-specific PLVs between the two AD and HC groups.

**Figure 2:**
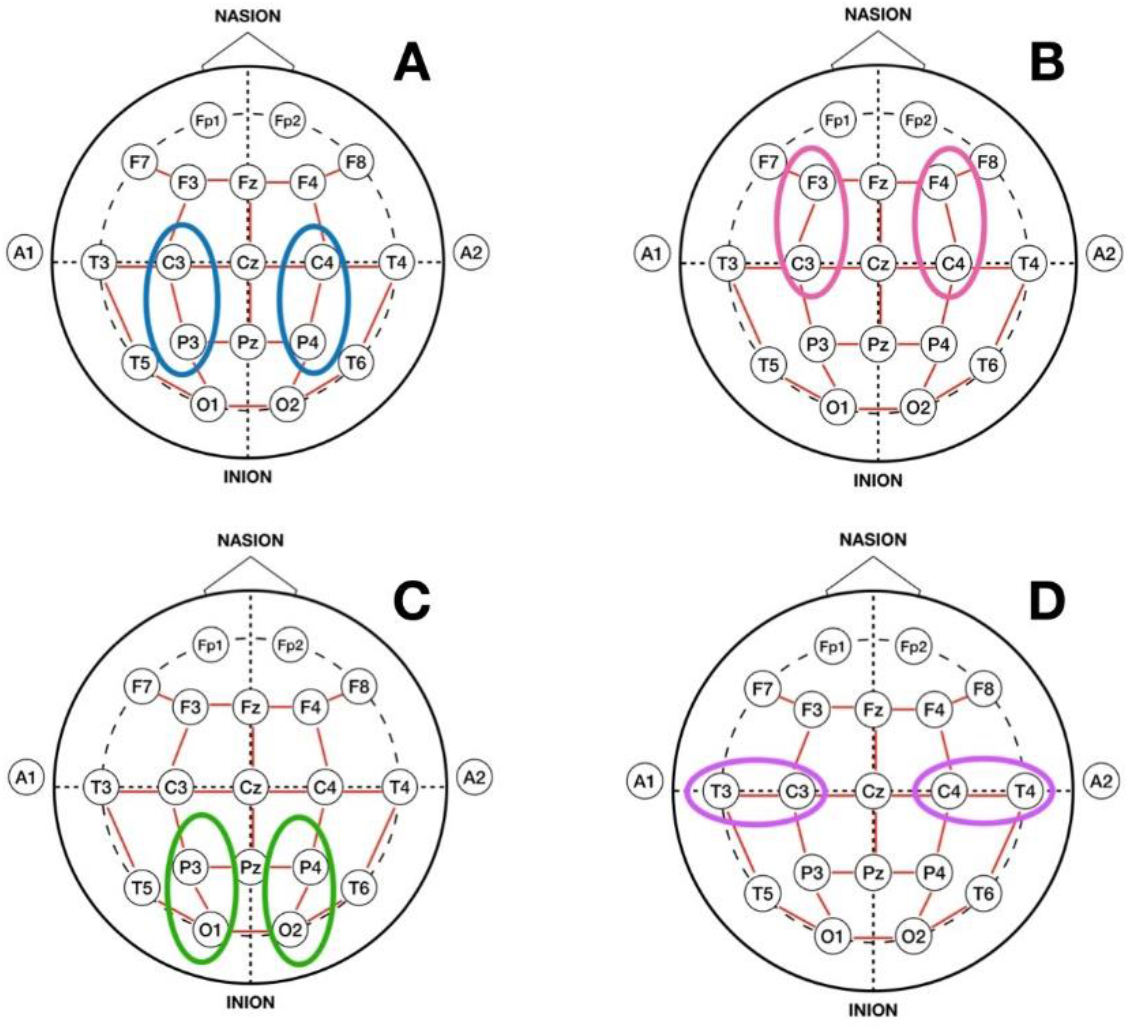
Homotopic pair connectivity was examined in four mirror regions in the left and right hemispheres including A) centro-parietal (C3-P3 & C4-P4), B) fronto-central area (F3-C3 & F4-C4), C) parieto-occipital (P3-O1 & P4-O2) and D) centro-temporal (C3-T3 & C4-T4) connections.

#### Localization of AD using Homotopic Pair Connectivity

To answer the secondary RQ, the four homotopic pairs were compared against each other to ascertain which areas displayed a significant connectivity difference between the two groups. To do this, the PLV values obtained from both subject groups in each of the above-mentioned homotopic pairs were compared using Linear Mixed Effects (LME) regression models. LME was fit using the lme4() package [32] and was chosen for this analysis because the repeated measure from the homotopic pairs were correlated, violating the assumptions of other tests, such as ANOVAs.

The analysis included two steps; first, the LME model was fit with PLVs as response variable and Pair and Group as predictors. Participants were included as a random factor in the model. The interaction term was included to prevent the overly enthusiastic outcome that there is a difference in connectivity between HC and AD for all pairs. Next, following verification of main effects, post-hoc comparisons were conducted between pairs to examine which brain regions showed significant difference between the two groups. These steps were only applied to the frequency bands that showed statistically significant difference between the AD and HC groups in at least one of the homotopic pairs in the ‘Homotopic Pair Connectivity’ analysis.

## Results

### Spectral Analysis

The Shapiro-Wilk test applied to the band power data rejected the null hypothesis of normal populations distributions (*p* < 0.05). Therefore, the non-parametric Mann-Whitney U-test was used to compare the groups in each frequency band. The test did not find any significantly different delta power for the AD subjects (*Mdn* = 4.23) than the healthy controls (*Mdn* = *4.07), U* = 174, *p* = 0.45. Similar results were observed for the theta (*Mdn* = 2.42 vs. *Mdn* = 4.30, *U* = 146, *p* = 0.15), alpha (*Mdn* = 1.88 vs. *Mdn* = 2.29, *U* = 162, *p* = 0.31), beta (*Mdn* = 1.67 vs. *Mdn* = 1.76, *U* = 152, *p* = 0.2) and gamma bands (*Mdn* = 0.67 vs. *Mdn* = 0.90, *U* = 158, *p* = 0.26). Therefore, it can be concluded that the spectral analysis yielded no significant differences between the AD subjects versus HC in any of the five frequency bands.

### Functional Connectivity Analysis

#### Global Connectivity

Figure 3 illustrates the distribution of PLV_mean_ from all subjects in the AD and HC groups in all five frequency bands. The result of the Mann-Whitney test indicated that the average PLVs from all channel combinations were significantly higher in the theta band for the AD participants (*Mdn* = 0.31) when compared to the HCs (*Mdn* = 0.26), *U* = 326, *p* = 0.0004. This was not the case for the delta (*Mdn* = 0.30 vs. *Mdn* = 0.28, *U* = 258, *p* = 0.12), alpha (*Mdn* = 0.26 vs. *Mdn* = 0.24, *U* = 264, *p* = 0.09), beta (*Mdn* = 0.18 vs. *Mdn* = 0.18, *U* = 226, *p* = 0.50) and gamma bands (*Mdn* = 0.18 vs. *Mdn* = 0.18, *U* = 181, *p* = 0.62).

**Figure 3:**
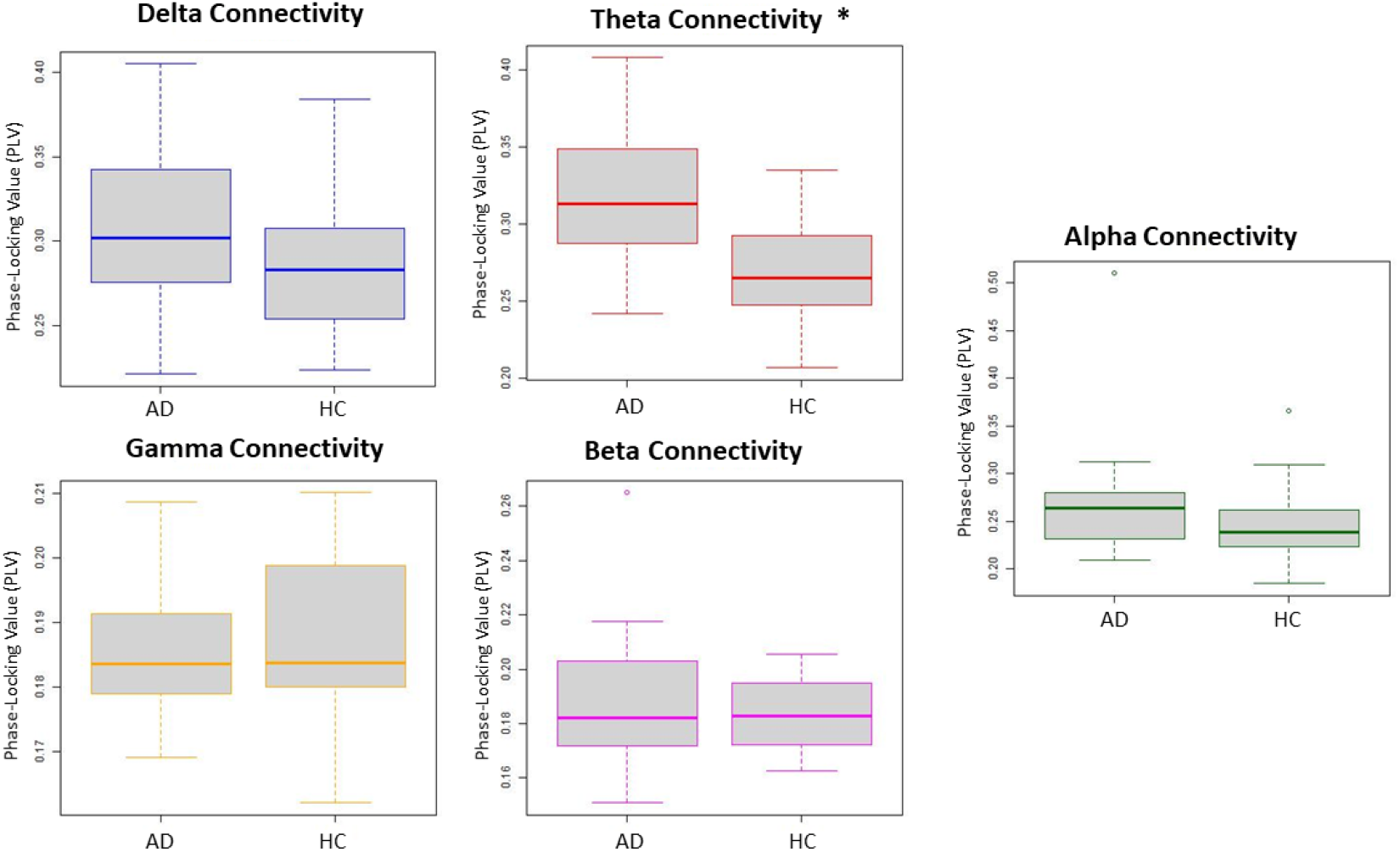
The average PLVs obtained from all connectivity pairs for the five frequency bands (Global Connectivity). Plots marked with * indicate statistically significant difference (*p* < 0.05) between AD patients and HCs.

#### Homotopic Pair Connectivity

Figure 4 illustrate the PLV values obtained from the homotopic pair in the centro-parietal area (Pair A) of the AD and HC groups in the five frequency bands. The Mann-Whitney test displayed a significantly higher PLV in the theta band for AD participants (*Mdn* = 0.64) as compared to HCs (*Mdn* = 0.52), *U* = 292, *p* = 0.01. No significant results were found for the other four frequency bands.

**Figure 4:**
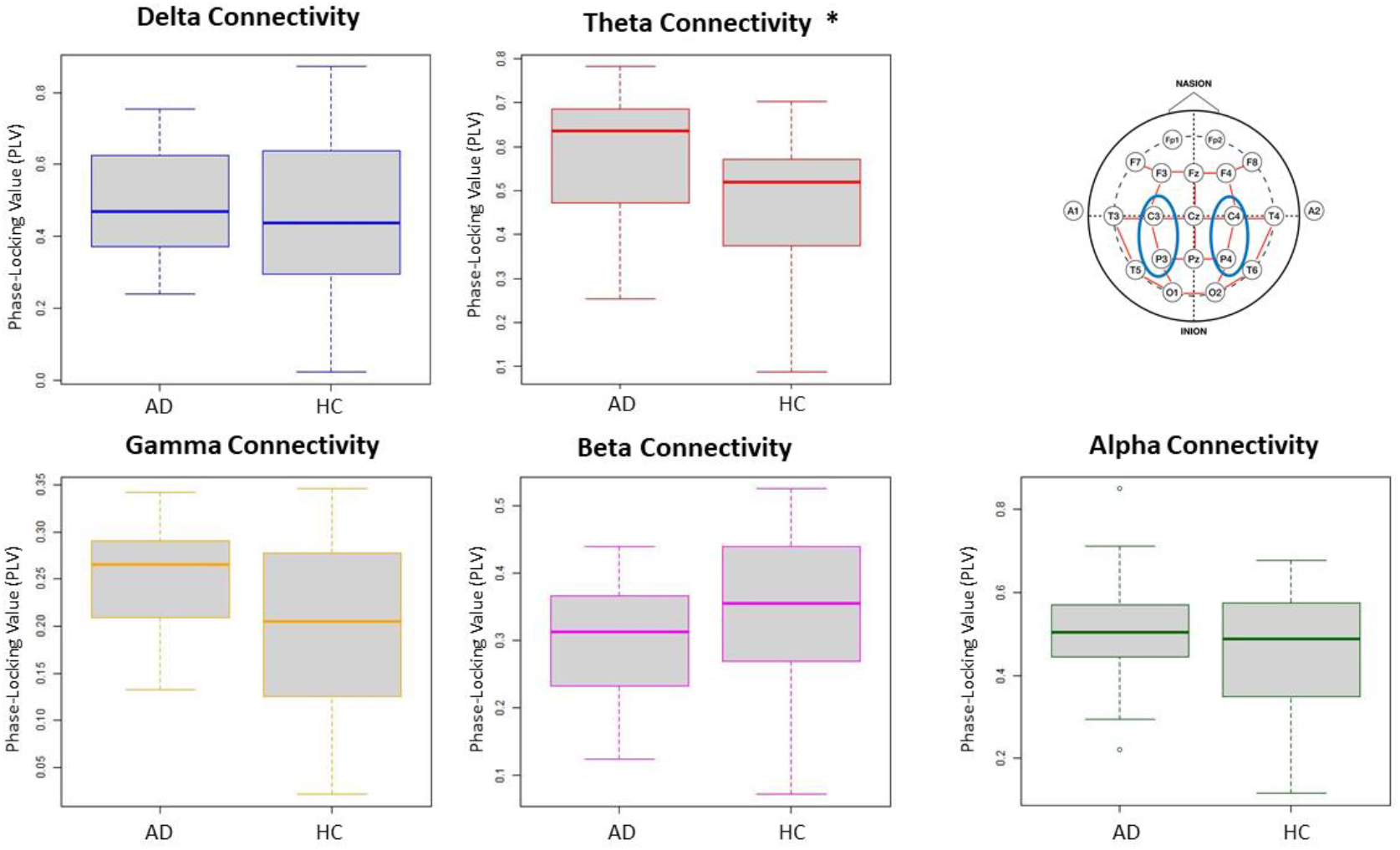
The PLVs obtained from the homotopic pair in the centro-parietal region (Pair A). Plots marked with * indicate statistically significant difference (*p* < 0.05) between AD patients and HCs.

Figure 5 illustrates the PLV values obtained from the homotopic pair in the fronto-central region (Pair B) of the AD and HC groups in the five frequency bands. The Mann-Whitney test displayed a significantly higher PLV for AD participants in both the delta (AD *Mdn* = 0.57, HC *Mdn* = 0.45, *U* = 275, *p* = 0.04) and theta bands (AD *Mdn* = 0.65, HC *Mdn* = 0.50, *U* = 282, *p* = 0.03). No significant results were found for the other three frequency bands.

**Figure 5:**
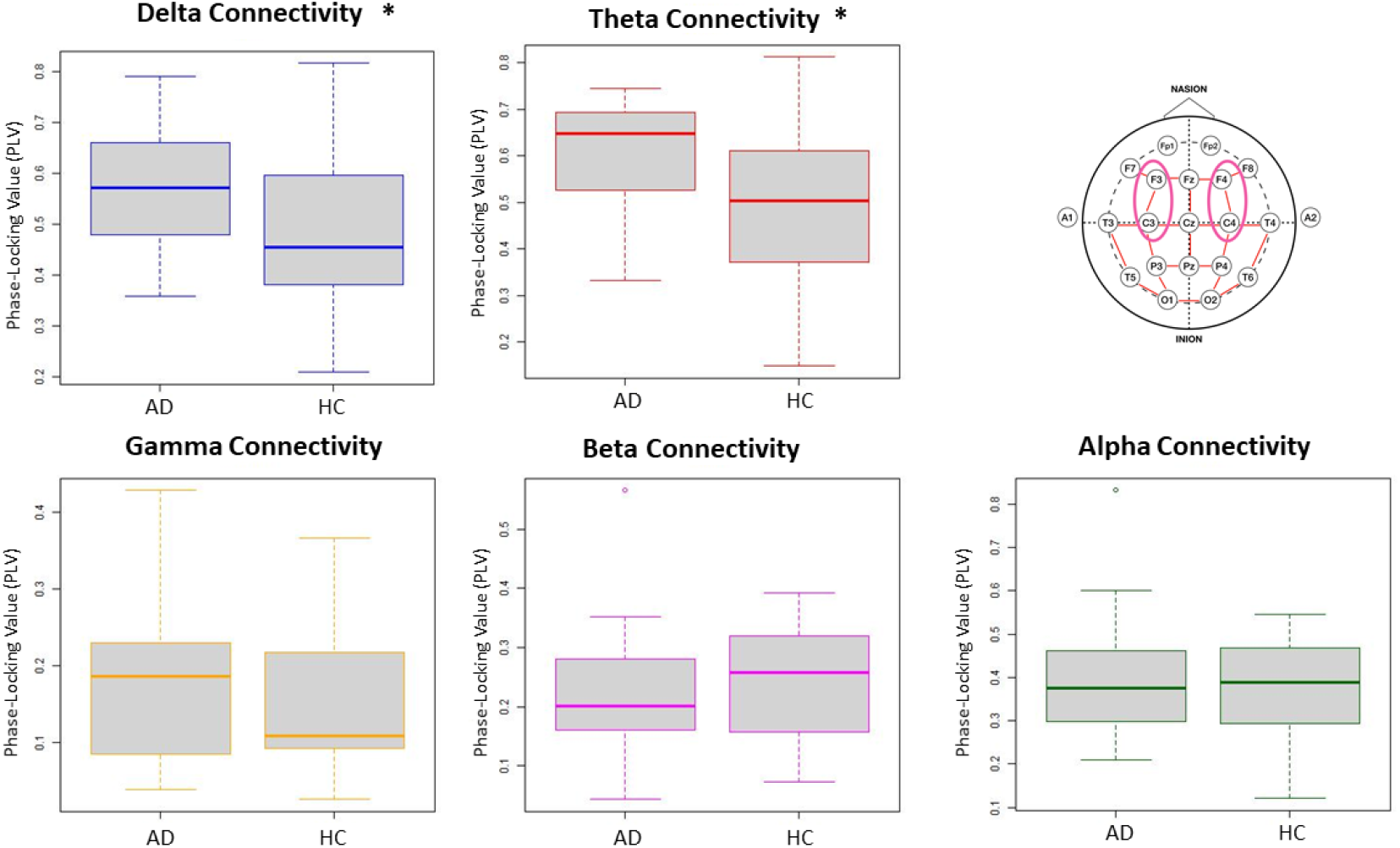
The PLVs obtained from the homotopic pair in the fronto-central region (Pair B). Plots marked with * indicate statistically significant difference (*p* < 0.05) between AD patients and HCs.

Figure 6 illustrates the PLV values obtained from homotopic pairs in the parieto-occipital region (Pair C) of the AD and HC groups in the five frequency bands. The Mann-Whitney test indicated a significantly higher PLV for the AD group (*Mdn* = 0.64) as compared to the HCs (*Mdn* = 0.49), solely in the delta band (*U* = 293, *p* = 0.01). The test resulted in insignificant outcome for the other four frequency bands.

**Figure 6:**
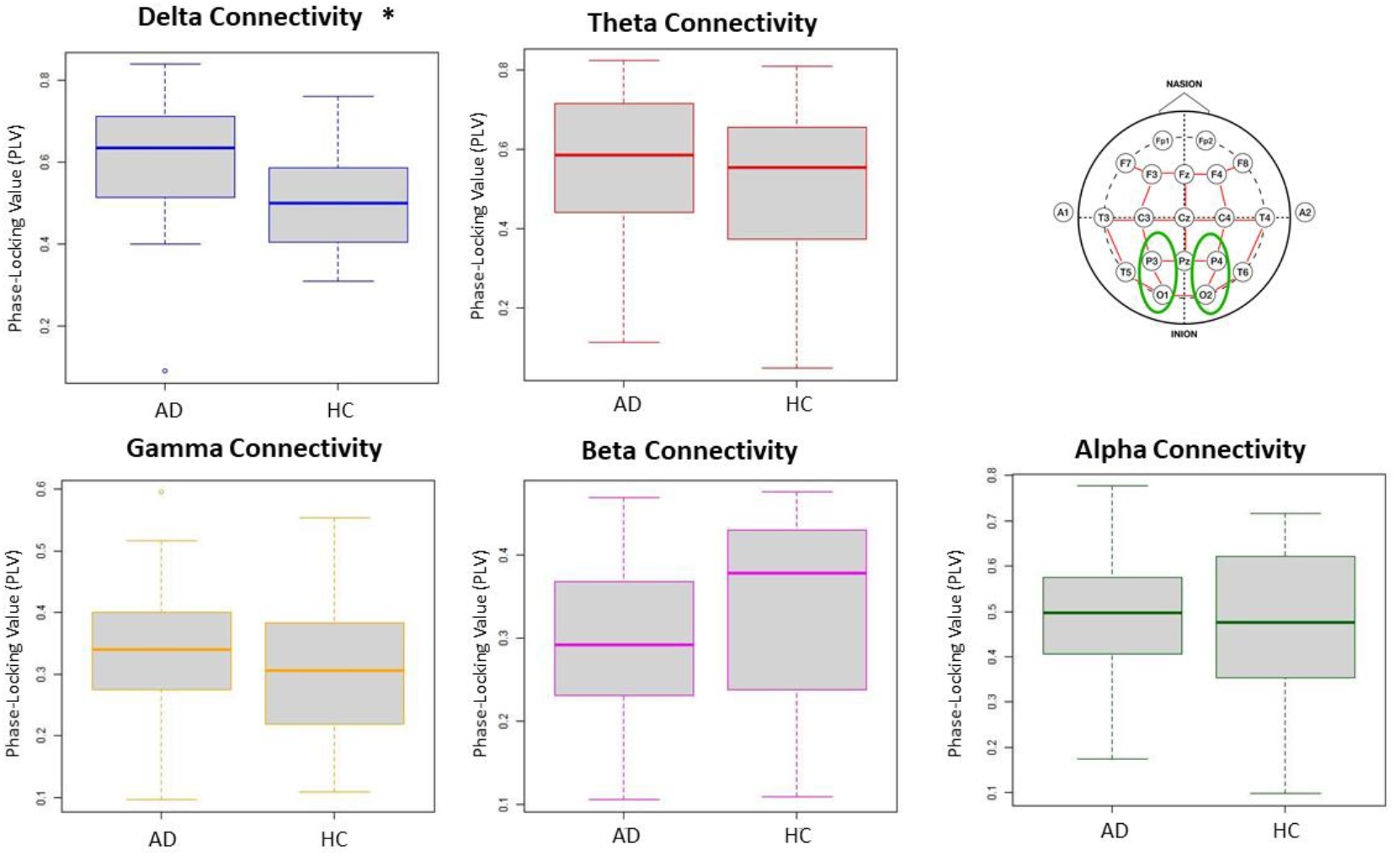
The PLVs obtained from the homotopic pair in the parieto-occipital region (Pair C). Plots marked with * indicate statistically significant difference (*p* < 0.05) between AD patients and HCs.

Lastly, Figure 7 shows the PLV values obtained from homotopic pairs in the centro-temporal region (Pair D) of the AD and HC groups in the five frequency bands. The Mann-Whitney test indicated a significantly higher PLV solely in the theta band of AD participants (*Mdn* = 0.48) as compared to the HCs (*Mdn* = 0.40, *U* = 280, *p* = 0.03). The results of group comparisons in the other four frequency bands remained insignificant.

**Figure 7:**
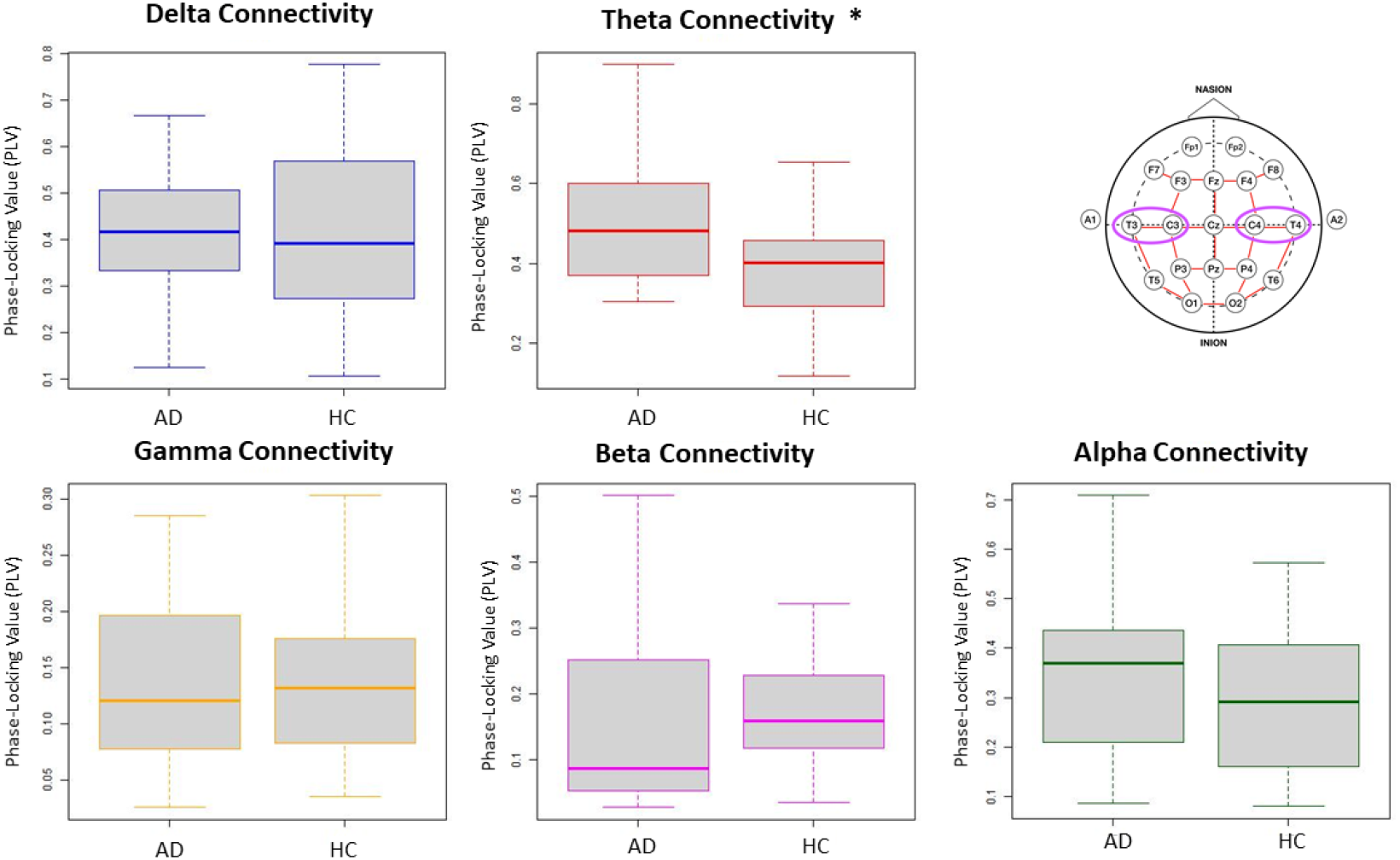
The PLVs obtained from the homotopic pair in the centro-temporal region (Pair D). Plots marked with * indicate statistically significant difference (*p* < 0.05) between AD patients and HCs.

In sum, the comparison of PLV in the selected homotopic pairs resulted in observing the main differences in the low frequency bands of delta and theta.

#### Localization of AD using Homotopic Pair Connectivity

To compare the connectivity across homotopic pairs and identify the most relevant brain region affected by AD, LME regression models were applied to the homotopic PLVs in the theta and delta frequency bands. The model confirming main effects for both Pair and Group was selected and post-hoc analysis using Tukey adjusted pairwise comparisons of least-squares means were conducted. Table 2 summarizes the outcome of post-hoc comparisons.

**Table 2:** Summary of the results of the post-hoc analysis of the LME regression.

| Homotopic Pair | Delta band (1 – 4 Hz) | Theta band (4 – 8 Hz) |
| --- | --- | --- |
| A | X | ✓ |
| B | X | X |
| C | X | X |
| D | X | ✓ |

In the theta band, the pairwise difference between AD patients and HCs reached significance for Pair A (*LSM difference* = 0.115, *SE* = 0.0517, *p* = 0.028) and Pair D (*LSM difference* = 0.101, *SE* = 0.0517, *p* = 0.038). While not statistically significant, trends were observed for Pair B (*LSM difference* = 0.097, *SE* = 0.0517, *p* = 0.064), whereas the difference between the AD and the HC group did not reach significance for Pair C. No significance was observed in the delta band.

## Discussion

The current study explored differences in the brain activity of patients afflicted with Alzheimer’s disease compared to a Healthy Control cohort, using quantitative analyses of EEG signals. In particular, two types of analyses were conducted and compared. First, a conventional spectral analysis was conducted to find spectral band power differences between AD subjects and healthy controls. The second approach employed a functional connectivity analysis using phase synchronization across five frequency bands to compare the intra-brain connectivity (global or local) between healthy brains and the ones affected by AD-induced dementia. The results indicated that the spectral analysis did not yield any significant differences between the AD and HC groups, suggesting that it is not an ideal method for diagnosis of AD based on EEG. On the other hand, the functional connectivity analysis using the PLV measure showed significant differences between the groups, both in terms of Global Connectivity and Homotopic Connectivity. Further analysis of homotopic pairs revealed significantly higher theta-band connectivity localized in the centro-parietal and centro-temporal regions.

The dataset used in this study consisted of bipolar derivations, instead of unipolar channel values that are more commonly used in the qEEG analysis [24, 29]. The use of bipolar derivations is seen as a more advantageous method compared to unipolar or average referencing methods [33], as it can mitigate the issues associated with common active referencing such as volume conduction [34]. Volume conduction, which refers to the leakage of electrical potentials to the neighboring electrodes, can complicate the interpretation of connectivity metrics. Therefore, the use of bipolar derivations in computation of functional connectivity is highly recommended as it was demonstrated in a recent study in the field of AD detection [30].

The results from the spectral analysis could not confirm any differences between the Alzheimer’s patients recruited in this study and their age-matched healthy controls. This is inconsistent with previous reports in which the development of AD was associated with an increase of delta and theta activity as well as a decrease in alpha and beta activity [6, 15]. An explanation for the lack of evidence in the current study could be that the AD subjects included in the sample were only moderately affected by this disorder. While this leaves room for future research to confirm the most suitable computation approach for detection of severe cases, this study proposes functional connectivity as a promising tool in detection of early signs of AD from EEG signals [35].

The two connectivity analyses that were subsequently carried out, namely ‘Global Connectivity’ and ‘Homotopic Pair Connectivity’, displayed increased communication between brain networks in the AD subject group when compared to the HCs. These findings were first identified in the Global Connectivity analysis and subsequently confirmed in the Homotopic Pair Connectivity analysis. The Global Connectivity analysis gave an overview of the AD process in the brain. Although it resulted in identifying higher connectivity distributed in the brains of AD group, it could not localize the effect. Indeed, the effect of Alzheimer’s disease tends to be more prominent in some areas of the brain than others [30, 31], hence justifying a motive to pursue a further analysis with the Homotopic Pair Connectivity. Similar to the band division performed to retrieve the PSD in a spectral analysis, in functional connectivity studies involving AD subjects, electrode pairs can be singled-out and evaluated separately, instead of combining them all together [36]. The analysis of homotopic pairs revealed a significant difference of connectivity in the delta band of the pairs in the fronto-central and the parieto-occipital regions whereas these effects were diminished in the Global Connectivity analysis which only found a significant difference between the groups in the theta band.

The result indicating a higher functional connectivity for the AD brains conflicts with the study of Hata et al. [10] who reported a lower lagged phase synchronization in delta and theta bands of AD patients. Indeed, a decreased connectivity between brain regions can be expected, as AD is known to cause neuronal loss and damage of neural pathways [1, 8, 20]. However, other studies suggest that the impact of such damage is only reflected on fast signals as healthy participants have higher brain connectivity in alpha and beta bands [17, 18] but not in the lower frequency bands. On the other hand, it has been shown in the past that patients suffering from neuropsychiatric disorders such as schizophrenia and epilepsy display increased functional activity between brain networks as a sign of anomaly in information communication [37, 38]. In the study of Cai et al. [15], similar patterns were reported for AD patients, where the connectivity within the same frequency band (intra-band connectivity) was stronger in AD brains than in the healthy brain whereas the connectivity between the frequency bands (inter-band connectivity) was significantly weaker. Observing higher synchronization values in the lower frequency bands for AD subjects can therefore be interpreted as a sign of brain dysfunction [15, 18]. More specifically, this study demonstrated that the brains affected by Alzheimer’s disease seemed to be in a ‘phase-lock’ state, causing a high connectivity in the low frequency bands; an observation that is well in line with the existing literature [15, 17, 30, 36, 39].

The “Localization of AD” analysis reached the conclusion that there was a significant difference in the connectivity between the AD and HC groups in the theta band for two out of four homotopic pairs. The answer to the secondary research question (RQ2) is therefore positive; it is possible to localize to some extent the differences between a healthy brain and one suffering from AD-induced dementia. The findings of this study therefore provide further evidence for damaged neural connections and consequently abnormal network dynamics in AD-affected brains particularly in the centro-parietal and centro-temporal regions. While older studies such as [40] suggested that the effects of AD are not situated in one specific area of the brain, the regions identified by this study are in line with the report of more recent studies such as Deng et al. [41] which observed a significant decrease in signal complexity of the AD group in the occipito-parietal and temporal regions of the brain using ‘multivariate multi-scale weighted permutation entropy’ (MMSWPE) [41].

Clearly, this study is not without limitations. A first limitation arises from the duration of the epochs that were available in the dataset (12s per subject). Longer epochs would have provided more EEG samples for phase synchronization analysis as well as an opportunity to evaluate the dynamic changes of connectivity over time as it had been previously done in Zhao et al. [30]. Another limitation involved the number of participants. The dataset used in this research consisted of 20 AD participants and 20 age-matched HCs. Given the individual differences inherent to the progress of AD, a larger dataset would have been optimal to yield more reliable results. Moreover, this study made use of the phase-locking value as an index for phase synchronization as the data was recorded in a bipolar manner and therefore the analysis was considered robust to the common source effects [27, 30]. Future research could use other indices of functional connectivity, such as coherence and phase-lag index (PLI), to investigate their efficacy is detecting AD impacts on the brain activity.

Finally, it shall be noted that this study applies a neural metric for Alzheimer’s detection from a data science perspective rather than a neuroscience one. This implies that the methodology employed in this study strived to find an accurate tool for detection of AD in EEG signals, rather than attempting to explain the cognitive and neural mechanisms the underlie the observed effects between AD participants and healthy controls. In this case, the findings of this research are well in line with the existing literature regarding AD detection and brain connectivity and show that the combination of bipolar derivations with phase synchronization can yield comparable results to studies that used other connectivity methods This qEEG analysis could therefore be considered as secondary tool, to be used alongside the visual EEG analysis employed by clinicians.

## Conclusion

This research served to find a promising tool for diagnosis of early signs of Alzheimer’s disease from brain activity by comparing two quantitative EEG methods, namely spectral analysis and functional connectivity analysis, in two groups of AD patients and age-matched Healthy Controls. The results indicated that the old-school spectral analysis failed to yield any statistically significant results that could help differentiate a brain affected by AD from a healthy one, whereas the functional connectivity analysis using phase synchronization found a significantly stronger global ‘phase-locked’ state in theta activity of AD-affected brains. Moreover, by extracting functional connectivity in four homotopic pairs of electrodes, it was possible to localize significant differences concerning the theta band in the centro-parietal and centro-temporal areas of the brain. To conclude, the findings of this research show that functional connectivity analysis using phase synchronization offers a promising quantitative method for future research in detection of AD. This method in combination with the standard cognitive tests that are commonly employed in dementia screening can put forward a more accurate diagnosis for patients who suffer from early symptoms of AD.

## Data Availability

The raw dataset used for this study is under a Non-Disclosure Agreement (NDA) and is therefore not available to the public.

The code used to support the findings of this study have been deposited in the GitHub repository (https://github.com/SemeliF/AD_paper).

## Acknowledgments

Authors would like to thank Dr. Ptolemaios G. Sarrigiannis from University of Sheffield for providing the EEG dataset used is this research, Sue Yoon from Eindhoven University of Technology for sharing her experience with the functional connectivity analysis and Dr. Peter Hendrix from Tilburg University for his guidance regarding the statistical analysis.

## References

[1] Neto, E., Allen, E. A., Aurlien, H., Nordby, H., & Eichele, T. (2015). EEG spectral features discriminate between Alzheimer’s and vascular dementia. Frontiers in Neurology, 6, 25.

[2] Smailovic, U. & Jelic, V. (2019). Neurophysiological markers of Alzheimer’s disease: Quantitative EEG approach. Neurology and Therapy, 8, 37 – 55.

[3] Wang, R., Wang, J., Yu, H., Wei, X., Yang, C., & Deng, B. (2014). Power spectral density and coherence analysis of Alzheimer’s EEG. Cognitive Neurodynamics, 9, 291–304.

[4] Heideman, M., Johnson, D., & Burrus, C. (1984). Gauss and the History of the Fast Fourier Transform. IEEE ASSP Magazine, 1(4), 14–21.

[5] Signorino, M., Pucci, E., Belardinelli, N., Nolfe, G., & Angeleri, F. (1995). EEG spectral analysis in vascular and Alzheimer dementia. Electroencephalography and Clinical Neurophysiology, 94(5), 313–325. https://doi.org/10.1016/0013-4694(94)00290-2

[6] Ponomareva, N. V., Selesneva, N. D., & Jarikov, G. A. (2003). EEG Alterations in Subjects at High Familial Risk for Alzheimer’s Disease. Neuropsychobiology, 48(3), 152–159. https://doi.org/10.1159/000073633

[7] Hampel, H., Lista, S., Teipel, S. J., Garaci, F., Nisticò, R., Blennow, K., Zetterberg, H., Bertram, L., Duyckaerts, C., Bakardjian, H., Drzezga, A., Colliot, O., Epelbaum, S., Broich, K., Lehéricy, S., Brice, A., Khachaturian, Z. S., Aisen, P. S., & Dubois, B. (2014). Perspective on future role of biological markers in clinical therapy trials of Alzheimer’s disease: A long-range point of view beyond 2020. Biochemical Pharmacology, 88(4), 426–449.

[8] Sperling, R., Dickerson, B., Pihlajamaki, M. M., Vannini, P., LaViolette, P., Vitolo, O., Hedden, T., Becker, J., Rentz, D., Selkoe, D., & Johnson, K. (2009). Functional Alterations in Memory Networks in Early Alzheimer’s Disease. NeuroMolecular Medicine, 12, 27–43.

[9] Leuchter, A. F., Cook, I. A., Newton, T. F., Dunkin, J., Walter, D. O., Rosenberg-Thompson, S., Lachenbruch, P. A., & Weiner, H. (1993). Regional differences in brain electrical activity in dementia: Use of spectral power and spectral ratio measures. Electroencephalography and Clinical Neurophysiology, 87(6), 385–393. https://doi.org/10.1016/0013-4694(93)90152-L

[10] Hata, M., Kazui, H., Tanaka, T., Ishii, R., Canuet, L., Pascual-Marqui, R. D., Aoki, Y., Ikeda, S., Kanemoto, H., Yoshiyama, K., Iwase, M., & Takeda, M. (2016). Functional connectivity assessed by resting state EEG correlates with cognitive decline of Alzheimer’s disease – an eLORETA study. Clinical Neurophysiology, 127(2), 1269–1278.

[11] Lombardi, A., Tangaro, S., Bellotti, R., Bertolino, A., Blasi, G., Pergola, G., Taurisano, P., & Guaragnella, C. (2017). A Novel Synchronization-Based Approach for Functional Connectivity Analysis. Complexity, 2017, 1–12.

[12] Babiloni, C., Lizio, R., Marzano, N., Capotosto, P., Soricelli, A., Triggiani, A. I., Cordone, S., Gesualdo, L., & Del Percio, C. (2016). Brain neural synchronization and functional coupling in Alzheimer’s disease as revealed by resting state EEG rhythms. International Journal of Psychophysiology, 103, 88–102.

[13] Fell, J. & Axmacher, N. (2011). The role of phase synchronization in memory processes. Nature Reviews Neuroscience, 12, 105–118.

[14] Bastos, A. M. & Schoffelen, J.-M. (2016). A Tutorial Review of Functional Connectivity Analysis Methods and Their Interpretational Pitfalls. Frontiers in Systems Neuroscience, 9, 175.

[15] Cai, L., Wei, X., Wang, J., Yu, H., Deng, B., & Wang, R. (2018). Reconstruction of functional brain network in Alzheimer’s disease via cross-frequency phase synchronization. Neurocomputing, 314, 490–500.

[16] Lachaux, J.-P., Rodriguez, E., Martinerie, J., & Varela, F. J. (1999). Measuring phase synchrony in brain signals. Human Brain Mapping, 8(4), 194–208.

[17] Yu, M., Gouw, A. A., Hillebrand, A., Tijms, B. M., Stam, C. J., van Straaten, E. C., & Pijnenburg, Y. A. (2016). Different functional connectivity and network topology in behavioral variant of frontotemporal dementia and Alzheimer’s disease: an EEG study. Neurobiology of Aging, 42, 150–162.

[18] Engels, M. M. A., Stam, C., van der Flier, W. M., Scheltens, P., de Waal, H., & van Straaten, E. V. (2015). Declining functional connectivity and changing hub locations in Alzheimer’s disease: an EEG study. BMC Neurology, 15.

[19] Bruña, R., Maestú, F., & Pereda, E. (2018). Phase locking value revisited: teaching new tricks to an old dog. Journal of Neural Engineering, 15(5), 056011.

[20] Poza, J., Bachiller, A., Gómez, C., García, M., Núñez, P., Gomez-Pilar, J., Tola-Arribas, M. A., Cano, M., & Hornero, R. (2017). Phase-amplitude coupling analysis of spontaneous EEG activity in Alzheimer’s disease. In 2017 39th Annual International Conference of the IEEE Engineering in Medicine and Biology Society (EMBC), (pp. 2259–2262).

[21] Schreiter-Gasser, U., Gasser, T., & Ziegler, P. (1994). Quantitative EEG analysis in early onset Alzheimer’s disease: correlations with severity, clinical characteristics, visual EEG and CCT. Electroencephalography and Clinical Neurophysiology, 90(4), 267–272. https://doi.org/10.1016/0013-4694(94)90144-9

[22] Gallego-Jutglà, E., Elgendi, M., Vialatte, F., Solé-Casals, J., Cichocki, A., Latchoumane, C., Jeong, J., & Dauwels, J. (2012). Diagnosis of Alzheimer’s disease from EEG by means of synchrony measures in optimized frequency bands. 2012 Annual International Conference of the IEEE Engineering in Medicine and Biology Society, 4266–4270. https://doi.org/10.1109/EMBC.2012.6346909

[23] Elgendi, M., Vialatte, F., Cichocki, A., Latchoumane, C., Jeong, J., & Dauwels, J. (2011). Optimization of EEG frequency bands for improved diagnosis of Alzheimer disease. Annual International Conference of the IEEE Engineering in Medicine and Biology Society. IEEE Engineering in Medicine and Biology Society. Annual International Conference, 2011, 6087–6091. https://doi.org/10.1109/IEMBS.2011.6091504

[24] Czigler, B., Csikós, D., Hidasi, Z., Anna Gaál, Z., Csibri, É., Kiss, É., Salacz, P., & Molnár, M. (2008). Quantitative EEG in early Alzheimer’s disease patients—Power spectrum and complexity features. International Journal of Psychophysiology, 68(1), 75–80. https://doi.org/10.1016/j.ijpsycho.2007.11.002

[25] Jasper, H. (1958). The Ten-Twenty Electrode System of the International Federation. Electroencephalography and Clinical Neurophysiology, 10, 371–375

[26] Rutkove, S. B. (2007). Introduction to Volume Conduction, (pp. 43–53). Totowa, NJ: Humana Press.

[27] Blackburn, D. J., Zhao, Y., De Marco, M., Bell, S. M., He, F., Wei, H.-L., Lawrence, S., Unwin, Z. C., Blyth, M., Angel, J., Baster, K., Farrow, T. F. D., Wilkinson, I. D., Billings, S. A., Venneri, A., & Sarrigiannis, P. G. (2018). A pilot study investigating a novel non-linear measure of eyes open versus eyes closed EEG synchronization in people with Alzheimer’s disease and healthy controls. Brain Sciences, 8(7), 134. https://doi.org/10.3390/brainsci8070134

[28] Yoon, S., Alimardani, M., & Hiraki, K. (2021). The Effect of Robot-Guided Meditation on Intra-Brain EEG Phase Synchronization. Companion of the 2021 ACM/IEEE International Conference on Human-Robot Interaction, 318–322. https://doi.org/10.1145/3434074.3447184.

[29] Leeuwis, N., Yoon, S., & Alimardani, M. (2021). Functional Connectivity Analysis in Motor-Imagery Brain Computer Interfaces. Frontiers in Human Neuroscience, 15, 564. https://doi.org/10.3389/fnhum.2021.732946

[30] Zhao, Y., Zhao, Y., Durongbhan, P., Chen, L., Liu, J., Billings, S. A., Zis, P., Unwin, Z. C., De Marco, M., Venneri, A., Blackburn, D. J., & Sarrigiannis, P. G. (2020). Imaging of Nonlinear and Dynamic Functional Brain Connectivity Based on EEG Recordings With the Application on the Diagnosis of Alzheimer’s Disease. IEEE transactions on medical imaging, 39(5), 1571–1581. https://doi.org/10.1109/TMI.2019.2953584

[31] Durongbhan, P., Zhao, Y., Chen, L., Zis, P., De Marco, M., Unwin, Z. C., Venneri, A., He, X., Li, S., Zhao, Y., Blackburn, D. J., & Sarrigiannis, P. G. (2019). A Dementia Classification Framework Using Frequency and Time-Frequency Features Based on EEG Signals. IEEE transactions on neural systems and rehabilitation engineering : a publication of the IEEE Engineering in Medicine and Biology Society, 27(5), 826–835. https://doi.org/10.1109/TNSRE.2019.2909100

[32] Bates, D., Mächler, M., Bolker, B., & Walker, S. (2015). Fitting Linear Mixed-Effects Models Using lme4. Journal of Statistical Software, 67(1), 1–48. https://doi.org/10.18637/jss.v067.i01

[33] Acharya, J. N., & Acharya, V. J. (2019). Overview of EEG Montages and Principles of Localization. Journal of Clinical Neurophysiology: Official Publication of the American Electroencephalographic Society, 36(5), 325–329. https://doi.org/10.1097/WNP.0000000000000538

[34] Trongnetrpunya, A., Nandi, B., Kang, D., Kocsis, B., Schroeder, C. E., & Ding, M. (2016). Assessing Granger Causality in Electrophysiological Data: Removing the Adverse Effects of Common Signals via Bipolar Derivations. Frontiers in Systems Neuroscience, 9, 189. https://doi.org/10.3389/fnsys.2015.00189

[35] Rossini, P. M., Di Iorio, R., Vecchio, F., Anfossi, M., Babiloni, C., Bozzali, M., Bruni, A. C., Cappa, S. F., Escudero, J., Fraga, F. J., Giannakopoulos, P., Guntekin, B., Logroscino, G., Marra, C., Miraglia, F., Panza, F., Tecchio, F., Pascual-Leone, A., & Dubois, B. (2020). Early diagnosis of Alzheimer’s disease: the role of biomarkers including advanced EEG signal analysis. Report from the IFCN-sponsored panel of experts. Clinical neurophysiology : official journal of the International Federation of Clinical Neurophysiology, 131(6), 1287–1310. https://doi.org/10.1016/j.clinph.2020.03.003

[36] Sankari, Z., Adeli, H., & Adeli, A. (2011). Intrahemispheric, interhemispheric, and distal EEG coherence in Alzheimer’s disease. Clinical neurophysiology : official journal of the International Federation of Clinical Neurophysiology, 122(5), 897–906. https://doi.org/10.1016/j.clinph.2010.09.008

[37] Yin, Z., Li, J., Zhang, Y., Ren, A., Meneen, K. M. V., & Huang, L. (2017). Functional brain network analysis of schizophrenic patients with positive and negative syndrome based on mutual information of EEG time series. Biomedical Signal Processing and Control, C(31), 331–338. https://doi.org/10.1016/j.bspc.2016.08.013

[38] Quraan, M. A., McCormick, C., Cohn, M., Valiante, T. A., & McAndrews, M. P. (2013). Altered Resting State Brain Dynamics in Temporal Lobe Epilepsy Can Be Observed in Spectral Power, Functional Connectivity and Graph Theory Metrics. PLOS ONE, 8(7), e68609. https://doi.org/10.1371/journal.pone.0068609

[39] Locatelli, T., Cursi, M., Liberati, D., Franceschi, M., & Comi, G. (1998). EEG coherence in Alzheimer’s disease. Electroencephalography and clinical neurophysiology, 106(3), 229–237. https://doi.org/10.1016/s0013-4694(97)00129-6

[40] Adeli, H., Ghosh-Dastidar, S., & Dadmehr, N. (2008). A spatio-temporal wavelet-chaos methodology for EEG-based diagnosis of Alzheimer’s disease. Neuroscience Letters, 444(2), 190–194. https://doi.org/10.1016/j.neulet.2008.08.008

[41] Deng, B., Cai, L., Li, S., Wang, R., Yu, H., Chen, Y., & Wang, J. (2017). Multivariate multi-scale weighted permutation entropy analysis of EEG complexity for Alzheimer’s disease. Cognitive Neurodynamics, 11(3), 217–231. https://doi.org/10.1007/s11571-016-9418-9

